# Increasing global heterogeneity in the life expectancy of older populations

**DOI:** 10.1101/589630

**Authors:** William Joe, Lathan Liou, S V Subramanian

**Author notes:** Both authors contributed equally (William Joe, Lathan Liou). Corresponding Author: SV Subramanian, Professor of Population Health and Geography, Harvard Center for Population and Development Studies, 9 Bow Street, Cambridge MA 02138.

## Abstract

With overall global improvements in life expectancy, one important concern is whether there is cross-country convergence in life expectancy at various ages. Insights in convergence patterns can help realign research priorities help governments better structure health investments across various age groups. We reveal global patterns in life expectancy improvements and identify convergent clubs in life expectancy at various ages for 201 countries / areas between 1950 and 2015. In the case of life expectancy at younger ages, most countries are moving in the same direction, but we observe significant cross-country variation for older adults and the elderly. Further, we observe increasing variance in life expectancy for older adults and elderly across countries. Increasing global heterogeneity in survival experience of older adults and the elderly population thus has remained a neglected aspect in the discussions on global life expectancy improvements. Data, research and policy focus beyond life-expectancy at birth is therefore critical to accelerate survival gains among older adults and elderly, particularly from the developing world.

## INTRODUCTION

The last two centuries witnessed unparalleled social and scientific advancements that have pushed back the frontiers of human longevity [1–3]. Global life expectancy at birth is estimated to have more than doubled in the past two centuries from 25 years to over 65 years [1, 35]. Given such improvements, there are important reasons for discerning global patterns of convergence in life expectancy at various ages. Because approaching convergence is intrinsic to achieving health equity, examining heterogeneities in life expectancy, particularly among older adults, is instrumental for realigning health and developmental priorities. Previously, life expectancy at birth has been a widely studied health metric, and studies have shown convergence in life expectancy at birth for a subset of industrialized countries [4–6]. However, an exclusive focus on life expectancy at birth may conflate early survival benefits with late survival gains in longevity. In fact, recent evidence from high-income countries indicate widening mortality differentials between older adults and the elderly population [33,36]. These insights warrant comprehensive assessment of cross-country patterns in life expectancy improvements across various age thresholds to reconcile a salient and emerging concern in global health.

This study examines global improvements in life expectancy at various ages and unravels patterns of convergence or its lack thereof. The notion of convergence is closely associated with health equity and development [37]. Previous studies on convergence discern significant improvements among countries with low initial life expectancies at birth [5,7,8]. This catch up potential among lagging countries coherently aligns with Omran’s theory of epidemiological transition [9]. The bulk of these improvements can be attributable to improving access to maternal and child health interventions as well as a reductions in the burden of infectious diseases afforded by widespread diffusion of cost-effective medical interventions [10]. While Wilson [5] was among the earliest to quantify the extent of global convergence, Mayer-Foulkes [11] observes that life expectancy improvements can be further explored through “convergence clubs”, which consist of countries displaying similar life expectancy dynamics. McMichael *et al*. [12], in fact, identified three distinct convergent clubs: those that have experienced rapid improvement, those that have experienced relative stagnation, and those that have experienced an erosion of life expectancy. Similarly, Bloom and Canning discuss a two-club model of convergence in life expectancy at birth whereby some countries are able to make the transition from an unhealthy, high-mortality club to a healthy, low-mortality club whereas others fail to escape the “mortality trap” [13]. But despite increasing relevance, there is a lack of research to unravel the global improvements and patterns of convergence in life expectancy at other ages beyond that at birth. This gap in evidence is partly attributable to data constraints and partly associated with a lack of methodological alternatives to test convergence and detect convergent clubs. Nevertheless, in recent years there have been progress on both fronts particularly with major improvements in availability of country-specific life expectancy estimates by UNDESA as well as robust methodological alternatives to test convergence.

This paper applies a novel econometric technique (the log-t convergence test) to examine convergence and detect convergence clubs and divergent group of countries [14]. The technical advantages of the method are two-fold [14]. First, this nonlinear time-varying regression allows for transitional divergence. This aligns with Vallin and Meslé, who emphasize on integrating divergence-convergence patterns that is expected with every advance in health since resourceful nations would benefit initially, causing divergence, until others can gradually access the health improvements, and subsequently converge [17]. Second, unlike previous studies [15,16], the method also allows for the method allows for identification of clusters that have similar life expectancy convergence / divergence patterns without any *a priori* assumptions.

We identify three distinct convergent clubs have emerged in life expectancy trends among the elderly population, which we are defining to be above the age threshold of 65. The life expectancy gains among older adults are also heterogeneous and the trajectories of each club are markedly different. These findings are consistent with emerging evidence on inequalities in quality of life between different regions of the world [13,15,16,18,19]. In fact, we observe decreasing variance at younger ages but increasing variance at older ages as postulated by the theory of epidemiological transition, particularly the shifting mortality burden from infants and young children to older adults. Survival in younger ages across countries is now much improved whereas longevity improvements in older ages are increasingly varied. Clearly, it is necessary to achieve rapid improvements in health outcomes of older adults and the elderly to sustain the momentum in global life expectancy improvements. This also implies that countries lagging in adult survival or those experiencing stagnation in adult mortality may require substantial social security and public health investments to further population longevity.

### Data and Methods

The analysis is based on the life expectancy estimates (both sexes aggregated) from the 2015 revision of the population estimates and projections by the Population Division, United Nations [20]. Although the revision was carried out for a total of 233 countries or areas, life expectancy estimates are only available for 201 countries or areas that had a population of 90,000 or more in 2015. The age-specific mortality estimates in the 2015 revision rely on multiple data sources and estimation methods (model-based, small area, direct and indirect estimates) based on data availability and reliability [20]. However, in several cases, particularly for the earliest period and for developing countries, these estimates are not internally consistent and require further adjustments. Standard demographic techniques are applied to smooth for under-enumeration and age-heaping as well as death distribution methods or the various combinations of age groups. Mortality rates at older ages are further validated against reference values using the Human Mortality Database (HMD). Also, for earlier periods and in countries with deficient old age mortality data, mortality rates at age 75 and over are smoothed based on the observed average rate of mortality increase by age available through HMD and Max Planck Institute for Demographic Research. Further, different estimates or rates are often noted for countries based on different data sources or analytical methods. These are, however, adjusted through robust trend-fitting based on available datasets. The 2015 revision is thus derived based on all data available (censuses, household surveys, vital and population registers, and analytical reports) to provide consistent peer-reviewed estimates for all countries / areas.

Summary statistics for life expectancies at various ages during 1950-55 to 2010-15 are presented. However, for brevity, we focus on eight different age, namely, e00, e01, e05, e25, e50, e65, e75, and e85, with life expectancy at birth and life expectancy at age 85 denoted by e00 and e85, respectively. As such, e00 is an important health metric summarizing overall survival experience of the population from birth. Life expectancy at age 1 (e01) and age 5 (e05) can offer insights on convergence status particularly when the child survives the risk of infant and under-five mortality. Divergence in life expectancy here would signify an early onset of variation in mortality experience across countries. Similarly, e25 is a critical marker of survival prospects of youth who are vulnerable to various risks related to communicable, non-communicable and accidents/injuries [38]. Life expectancy at age 50 (e50) is an important age threshold for older adults and has been increasingly focused in global health [39]. A divergence here may reveal differentials in survival experience on account of elevated risk of NCDs. The focus on e65 is customary (standard international age threshold to define an elderly person) and depicts survival experience of the elderly population. Finally, e75 and e85 are used to examine whether there is increasing longevity gains for the population as a whole and whether it varies across countries [1]. Nevertheless, for completeness, we also present econometric analysis for all the available age cut-offs in the supplementary appendix.

Further, we examine variations in life expectancies improvements using multilevel linear regression models. The multilevel model is composed of three levels: year (level 1), country (level 2), and geographic region (level 3), which is classified according to 2015 Revision. We first use a random intercepts model to explore patterns of between-region and between-country variations in life expectancy at different age thresholds. More formally, we specify the following model:

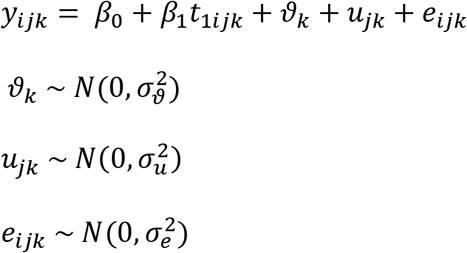

where, y_ijk_ is the life expectancy for a given age threshold in year i for country / area j and geographical region k, β_0_ is the mean life expectancy across all countries and time periods, β_1_t_1ijk_ is the slope coefficient for time variable introduced at level 1, ν_k_ is the effect of school k, u_jk_ is the effect of country / area within school region k, and e_ijk_ is the residual error term. The random effects and residual errors are assumed to be uncorrelated and normally distributed with zero means and constant variances. The variance parameters are used to compute the variance partition coefficient (VPC), and summarize the proportion of the total accounted variance in life expectancy at any specific level. The level-specific VPC is computed as the ratio of the level-specific variance (ν_k_ or u_jk_) to the total variance (ν_k_ + u_jk_ + e_ijk_).

Further, we also use the following two-level random slopes model to examine how between-country variance has evolved over the years for life expectancy at different age thresholds.

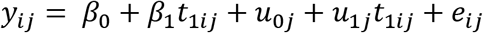

In this model, the country-level variance is the function of time and is computed following Leckie & Charlton [23]. In addition, the slope-intercept covariance from a similar three-level random slopes model is estimated to understand whether countries and regions with higher initial life expectancy experiences slower than average increments. In particular, a significantly large and negative covariance would offer preliminary insights on potential cross-country convergence.

Finally, we employ the log t convergence testing [14,21] to discern whether there is an overall convergence in life expectancies at various age thresholds. This approach models variance in life expectancy over time and estimates a relative transition coefficient to capture the life expectancy trajectory of a particular country relative to the cross-country average. Based on the coefficient, the trajectories of each country vis-à-vis the common trend is visualized using relative transition paths and divergent patterns are identified. Further, a time-series specification based on the relative transition parameters is used alongside a one-sided t-test to confirm acceptance of the null hypothesis of convergence vis-à-vis alternatives of no convergence or club convergence. Further details pertaining to the log-t test are presented in the supplementary appendix. The log-t test is performed using *psecta* module [22] whereas the multilevel analysis is performed using *runmlwin* module in Stata 15.0 [23].

## RESULTS

### Descriptive Analysis

Figure 1 confirms global improvements in life expectancy at various ages since the 1950s (Figure 1) [24–26]. The median lines in the box plot, however, also highlights a deceleration in life expectancy gains over time at e00, e01, e05, e25, and e50 [27]. Also notable is the increase in spread (inter-quartile range) of life expectancy at older ages. More clearly from Figure S1, we observe that variance in life expectancy across countries has a decreasing trend between the e00 and e25 age points but an increasing trend between the e50 and e85 age points. This hints that life expectancies at younger ages is shrinking whereas there is increasing variance at older ages.

**Figure 1:**
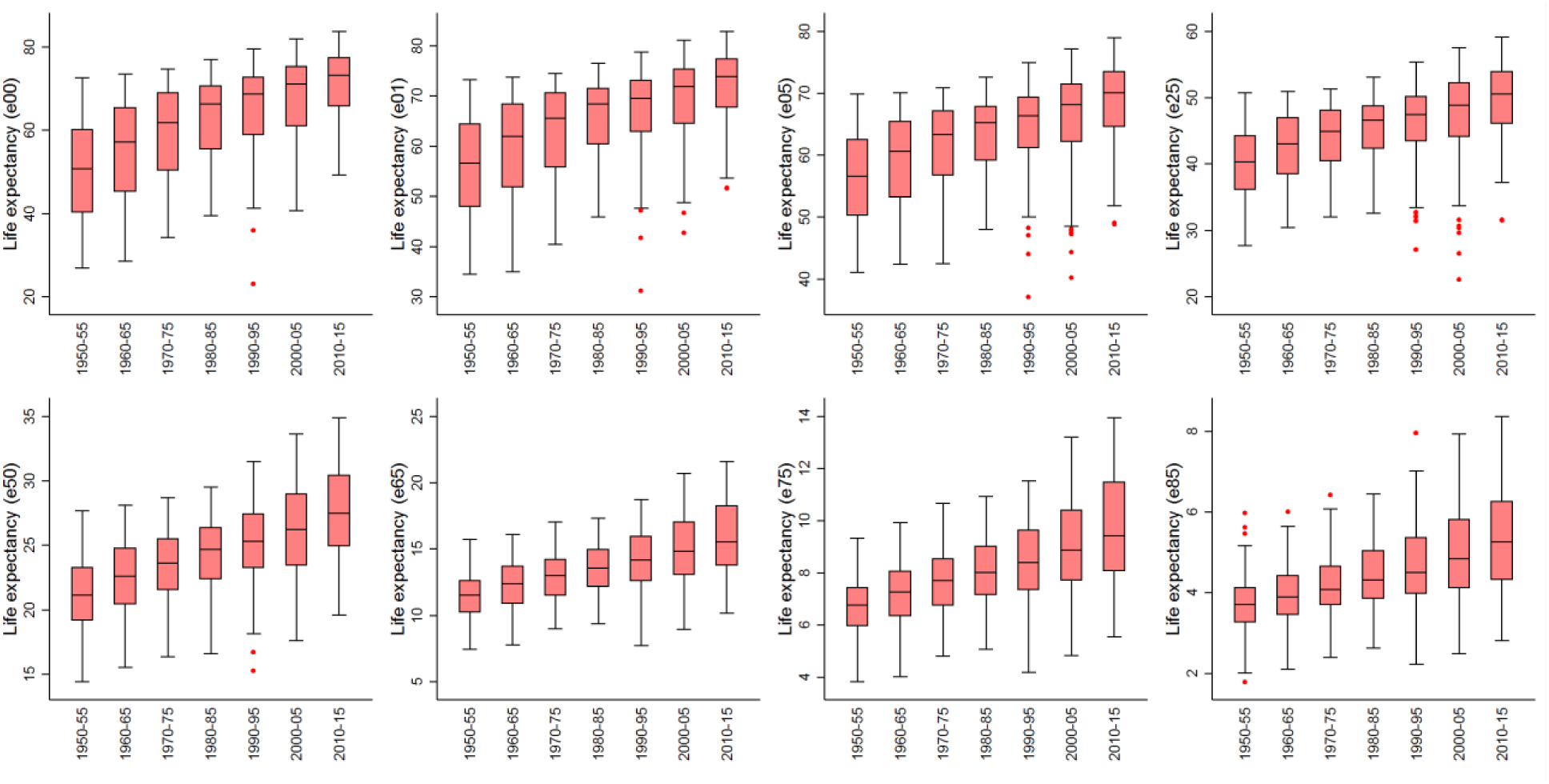
Box plot for life expectancy across countries at various ages, 1950-55 to 2010-15

The association between changes in life expectancy at birth vis-à-vis the base level (1950-55) indicates that larger gains have accrued to countries that had lower initial levels (Figure S2). Nevertheless, the pattern of base-level advantage dilutes when life expectancy gains at higher ages are considered. In particular, change in life expectancy at age 75 provides no advantage to countries with lower base levels. This can be due to slow progress among countries with low life expectancy or sustained increments among better-performing countries or both. These patterns are also indicative of socioeconomic, behavioral and other structural factors that influence survival of the older adults and the elderly [13,16].

### Multilevel Regression

We also perform multilevel regressions to understand the contributing factors to the life expectancy trends we observe. The random intercepts model estimates that geographic region accounts for 58.6% of the total variance of life expectancy at birth, whereas country and year account for 29.4% and 12.0% respectively (Table S1). At the other end of the age spectrum geographic region, country and year accounts for 40.0%, 37.4%, and 22.6% respectively of variance in life expectancy at age 85. Interestingly, at older ages, region seems to be relatively less heterogeneous whereas the between-country variations are rather salient. Further, the slope-intercept covariances from the random slopes model are significantly negative particularly at younger ages thus indicating potential for convergence whereas such no such association is inferred at the older ages (Table S2). The slope-intercept covariance at e00 is −0.535 [95% CI: −0.682; −0.389] and systematically decreases across the age spectrum. It is estimated to be −0.035 [95% CI: −0.048; −0.022] at e50 and further diminishes to −0.013 [95% CI: −0.019; −0.007].

### Convergence Analysis

The relative transition parameters during 1950-2015 is presented in Figure 2 to visualize the life expectancy trajectory of a particular country relative to the cross-country average. In the case of life expectancy at birth (e00), the relative transition curves across countries are moving towards constancy along with a clear shrinkage in heterogeneity across countries. However, a similar pattern of reduced dispersion cannot be discerned for the relative transition curves for life expectancy among older adults and elderly. In fact, a few countries such as Lesotho, Mali and Sierra Leone are found to be diverging from the pack.

**Figure 2:**
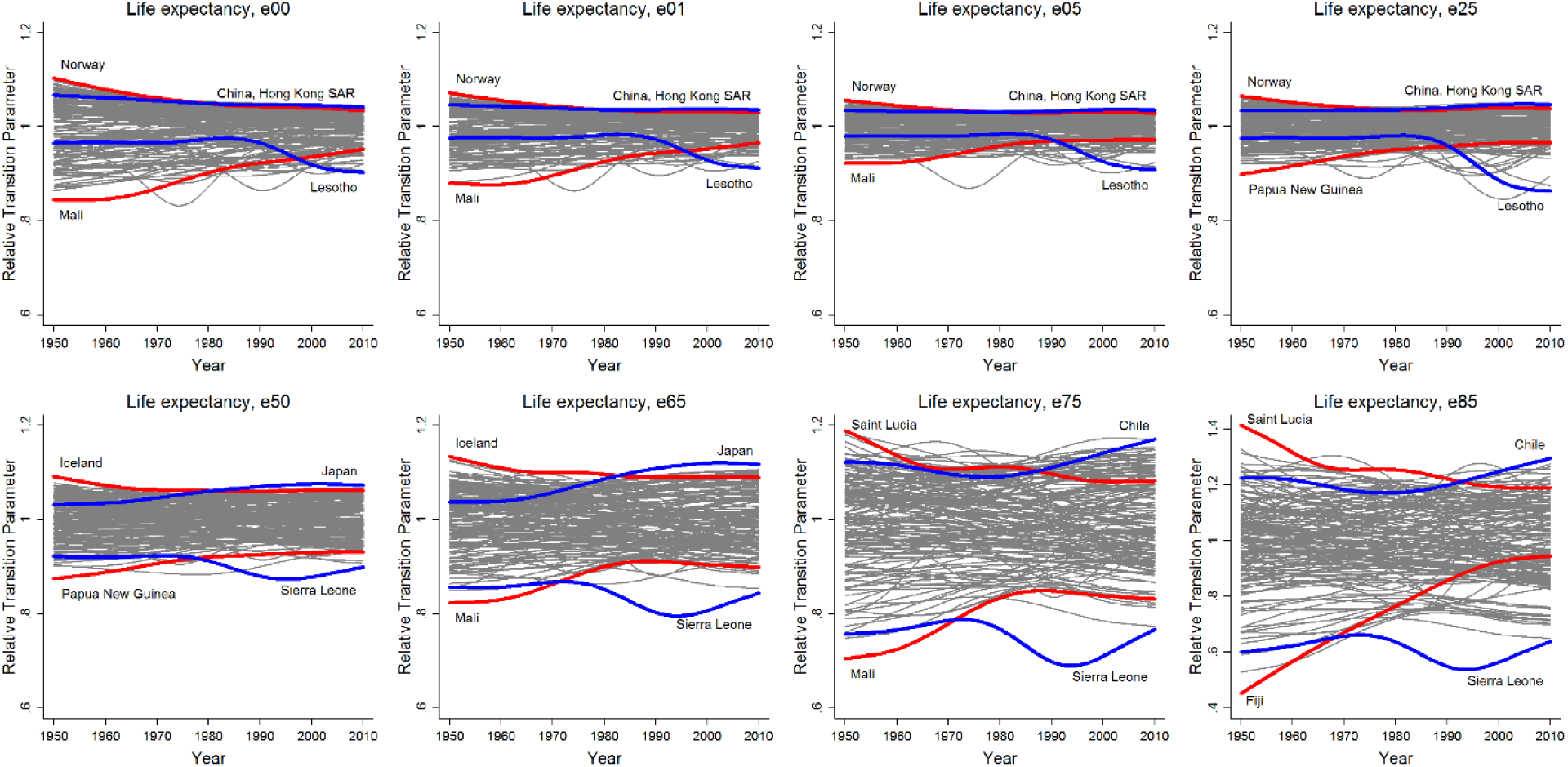
Country transition paths in life expectancy relative to world average, 1950-55 to 2010-15

Table 1 reports the *log-t* convergence test results with the corresponding club allocations at each age threshold (Supplemental Table S3). The results confirm that only in case of life expectancy at birth (γ = 0.202, 95% CI: 0.274; 0.131) are all the countries found to be converging whereas grand convergence is not observed for life expectancies at any other age. Nevertheless, in the case of life expectancies at younger ages (e01, e05, e25), most of the countries (over 170) are found to be part of one large common club but overall convergence fails because a few countries are divergent, notably Lesotho and Swaziland. In the “high mortality” convergence club are African countries, former Soviet Union countries, and countries that have in recent decades been subjected to immense political turmoil or war. In fact, the collapse of the Soviet Union has had adverse impact in health and life expectancy in Eastern Europe [15,16,19], and these results are coherent with previous findings. In older age cohorts, particularly former Soviet Union countries diverge from the other countries and are found to converge in another club. These seem to represent the countries that are making the transition to a higher-mortality cluster. The result is that in the e75 and e85 age groups, the club membership is rather balanced. For instance, for life expectancy at age 75, the countries can be divided into three broad clubs containing 61 (γ = 0.162, 95% CI: 0.274; 0.131), 62 (γ = 1.069, 95% CI: 1.334; 0.804) and 78 (γ = 0.148, 95% CI: 0.184; 0.112) countries. A complete list of countries and club membership is presented in Supplemental Table S4.

**Table 1:**
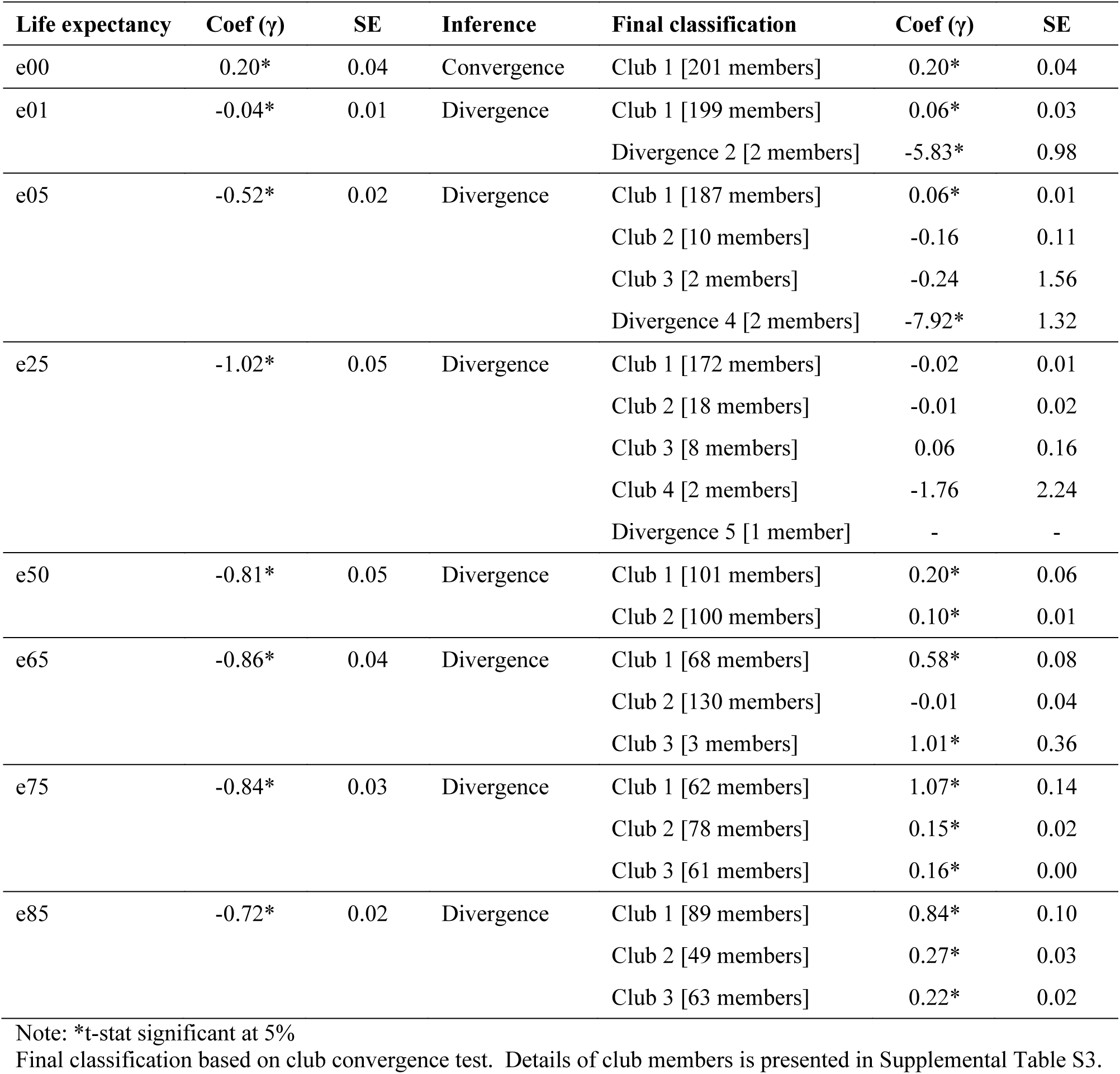
The Log-t test for convergence, club convergence and divergence in life expectancy at various ages across countries, 1950-55 to 2010-15

| Life expectancy | Coef ($\gamma$) | SE | Inference | Final classification | Coef ($\gamma$) | SE |
| --- | --- | --- | --- | --- | --- | --- |
| e00 | 0.20* | 0.04 | Convergence | Club 1 [201 members] | 0.20* | 0.04 |
| e01 | -0.04* | 0.01 | Divergence | Club 1 [199 members] | 0.06* | 0.03 |
|  |  |  |  | Divergence 2 [2 members] | -5.83* | 0.98 |
| e05 | -0.52* | 0.02 | Divergence | Club 1 [187 members] | 0.06* | 0.01 |
|  |  |  |  | Club 2 [10 members] | -0.16 | 0.11 |
|  |  |  |  | Club 3 [2 members] | -0.24 | 1.56 |
|  |  |  |  | Divergence 4 [2 members] | -7.92* | 1.32 |
| e25 | -1.02* | 0.05 | Divergence | Club 1 [172 members] | -0.02 | 0.01 |
|  |  |  |  | Club 2 [18 members] | -0.01 | 0.02 |
|  |  |  |  | Club 3 [8 members] | 0.06 | 0.16 |
|  |  |  |  | Club 4 [2 members] | -1.76 | 2.24 |
|  |  |  |  | Divergence 5 [1 member] | - | - |
| e50 | -0.81* | 0.05 | Divergence | Club 1 [101 members] | 0.20* | 0.06 |
|  |  |  |  | Club 2 [100 members] | 0.10* | 0.01 |
| e65 | -0.86* | 0.04 | Divergence | Club 1 [68 members] | 0.58* | 0.08 |
|  |  |  |  | Club 2 [130 members] | -0.01 | 0.04 |
|  |  |  |  | Club 3 [3 members] | 1.01* | 0.36 |
| e75 | -0.84* | 0.03 | Divergence | Club 1 [62 members] | 1.07* | 0.14 |
|  |  |  |  | Club 2 [78 members] | 0.15* | 0.02 |
|  |  |  |  | Club 3 [61 members] | 0.16* | 0.00 |
| e85 | -0.72* | 0.02 | Divergence | Club 1 [89 members] | 0.84* | 0.10 |
|  |  |  |  | Club 2 [49 members] | 0.27* | 0.03 |
|  |  |  |  | Club 3 [63 members] | 0.22* | 0.02 |

Note: \*t-stat significant at 5%
Final classification based on club convergence test. Details of club members is presented in Supplemental Table S3.

## DISCUSSION

We have illustrated that the age-aggregated life expectancy has masked the true pattern of stagnated gains in life expectancy for older populations. In particular, change in life expectancy at older ages continues to favor countries with higher life expectancy. Both high-income and low-income countries showed similar magnitudes in life expectancy gains. While poor performing countries are catching up with better performing ones, such sustained increase in life-expectancy improvements at older ages among better-performing countries is associated with continuing advancements in medical sciences as well as general improvements in health behaviors as well as improving physical and financial accessibility to advanced health care [6,10,17,27,28,30,39]. In addition, we observed that variance in life expectancy across countries has a decreasing trend between the e00 and e25 age points but an increasing trend between the e50 and e85 age points. This implies that the inequalities in life expectancy are declining at lower ages and increasing more quickly for the older populations. This finding coupled with an earlier observation regarding increasing between-country variance in life expectancy at older ages, assumes salience and calls for a renewed emphasis on between-country disparities in life expectancy in older populations. To a large extent, the increasing between-country variance has been attributed to major political turmoil (such as the collapse of communist regimes in Central Europe and Eastern Europe), the AIDS epidemic as well as varying pace of diffusion as well as access to medical technology [6].

There are two distinct convergent clubs for life expectancy at age 50 (e50 threshold), which resonates with Bloom and Canning’s high mortality - low mortality dual regime model. However, we show in the older age cohorts that there is a distinct three club pattern that better captures the nuances in mortality trajectories between countries. Countries without the means to achieve improved rates of diffusion of health technologies and implementation of public health measures [31,32] face a discouraging outlook of being stuck in a mortality trap [13]. In fact, older adults in low-income countries are particularly at higher risk of persistent mortality trends driven by non-communicable diseases [33]. The convergence clustering analysis confirms this persistent mortality trap thesis by identifying club of countries that are not performing very well with respect to life expectancy improvements specifically among the elderly (e65, e75, e85). Although it is promising that we have witnessed absolute life expectancy gains, it is clear that in order to achieve further increments in life expectancy, we have to reduce the mortality risks of the older adults stuck in the mortality trap. In this regard, cost-effective health interventions such as promoting responsible consumption behavior and curbing avoidable health risks such as tobacco use and unhealthy life-style and dietary practices can render substantive impact on life expectancy at older ages. In fact, efforts for mainstreaming preventive mechanisms are still in its infancy, particularly across the low-income countries [10]. It is important that these changes correspond with the epidemiological burden as sole reliance on increasing health care expenditures does not necessarily lead to convergence in health outcomes [18].

Finally, we consider the limitations and strengths of the paper. First, we used UNDESA 2015 Revisions, which is based on all data available including censuses, surveys, vital registers, international databases and specific modeling assumptions [20]. Although 2015 Revisions are consistent and have been peer-reviewed, we acknowledge that given the lack of quality data for certain countries especially in the 1950s, there remains uncertainty in modeling life expectancy for a number of developing countries. Nevertheless, our convergence results are still supported by previous research that also preclude convergence in life expectancy at other ages [2,4,5,8,25,34]. Second, this analysis does not examine the specific causes of our findings. Further research can thus explore life expectancy variations vis-à-vis causes of death as well as social determinants such as gender, income, and education.

In conclusion, we reiterate the need for accelerating survival gains among the older adults and the elderly, particularly in the developing world. Alternative survival indicators such as life expectancy at age 50 and age 65 should be routinely used to monitor survival disparities and health inequalities among the older adults. A continued improvement in life expectancy of the elderly population in the developed world invariably highlights the impact of socioeconomic transformation in enhancing longevity prospects of the population. But at the same time, countries experiencing sociopolitical instability, (particularly the African nations, Eastern European countries, and South Asian countries) devoid of certain fairly basic entitlements of health and livelihood is inimical to human rights and intensifies disparities in adult and elderly survival. Notwithstanding the need for sustaining improvements in maternal and child health, simultaneous efforts and impetus on health and well-being of older adults and the elderly is a growing necessity as humanity continues to push back the frontiers in human longevity.

## Author Contributions

W. Joe, L. Liou, and SV Subramanian equally contributed to the conceptualization and design of the study. W. Joe and L. Liou led the analysis and interpretation of data, co-drafted the initial manuscript, and reviewed the manuscript for important intellectual content. S.V. Subramanian provided overall supervision.

